# Serotype and distribution of adhesion genes in *Streptococcus mutans* clinical isolates

**DOI:** 10.1101/2024.04.09.588668

**Authors:** Laura Donnet, Olivier Claisse, Johan Samot

## Abstract

The delicate balance of the oral cavity is disrupted by hygiene and diet changes, leading to caries. The virulence of *Streptococcus mutans*, a key caries pathogen, is partly explained by its rhamnose-glucose polysaccharide (RGP) and its sucrose-dependent and -independent adhesion mechanisms. This study investigates the diversity of *S. mutans* through analysis of its RGP and the distribution of sucrose-independent adhesion proteins (SpaP, WapA, Cnm, Cbm) in Parkinson’s disease patients.

In the PARKIDENT clinical trial, strains were isolated from saliva samples of Parkinson’s patients before and after oral hygiene procedures. Strains were preanalyzed by multiplex PCR for serotype and collagen-binding proteins, followed by high-throughput sequencing and bioinformatics analysis using RAST® and BLAST®. Phylogenetic analyses and protein modeling were also conducted.

Of 40 patients in the Parkident clinical trial, only 24 had *Streptococcus mutans* strains isolable from salivary swabs. Serotyping revealed that over 80% of the 44 strains isolated were serotype c, in line with prevalence data in the literature. Furthermore, only SpaP types A and B were identified in the strains. The low variability observed for the WapA protein underlines its functional importance. Finally, the *cbm* gene was not found in any strain, and only 5 strains possessed the *cnm* gene, all of them serotype c.

The study of these 44 *Streptococcus mutans* strains showed characteristics similar to other populations. Serotype c predominates, with minimal peptide variability in adhesion proteins (SpaP, WapA). Collagen-binding proteins Cnm and Cbm are rare or absent. Further research is essential for detailed strain distribution in Parkinson’s-affected and general French populations due to *S. mutans’* growing role in extra-oral pathologies.

## Introduction

The oral cavity is a complex ecosystem in which a delicate balance is maintained. Among other factors, slight but repeated or continuous hygienic and dietary modifications will upset this balance. The consequence is dysbiosis, which can be the starting point for infectious pathologies of the mouth. Among these pathologies, caries remains a global public health challenge (Collaborators et al., 2020; Selwitz et al., 2007). Despite the involvement of other micro-organisms, *Streptococcus mutans* remains, for many authors, a key pathogen in the initiation of carious lesions (Takahashi & Nyvad, 2011; Tanner et al., 2018; Tanzer et al., 2001). This bacterium has adapted remarkably well to its environment and persists in the mouth via two principal adhesion mechanisms (Krzysciak et al., 2014). A sucrose-dependent adhesion mediated by exopolysaccharides synthesized by glucosyltransferases (Gtf B, C and D): these exopolysaccharides together with the expression of different glucan-binding proteins (Gbps) enable the bacterium to adhere to the hard tissues of the tooth and form part of the dental biofilm, which they help to stabilize and grow (Bowen & Koo, 2011; Matsumi et al., 2015). The other, a non-sucrose-dependent adhesion mechanism that involves surface adhesins such as SpaP, WapA and the collagen-binding adhesins Cnm and Cbm (Aviles-Reyes et al., 2017). SpaP (also known as P1, Pac, Antigen I/II) is recognized for its essential role in the initial adhesion of *S. mutans* to the tooth, as it binds to the product of the DMBT1 (Deleted in Malignant Brain Tumor 1) gene: salivary agglutinin (SAG), also known as SALivary Scavenger and Agglutinin (SALSA) or glycoprotein 340 (gp340). Three variants (A, B and C) of the SpaP protein have been described to date (Esberg et al., 2017). WapA is also known to participate in initial adhesion by binding to the exogenously acquired salivary pellicle covering the hard tissues of the tooth (Zhu et al., 2006). Both proteins have been shown to bind to collagen-rich surfaces (Aviles-Reyes et al., 2017; Beg et al., 2002).

This is also the case for the Cnm and Cbm adhesins, which, unlike the above proteins, are inconstantly present in the *S. mutans* genome. Their presence could amplify the pathogenic potential of *S. mutans*, as several authors have reported their presence in strains implicated in extra-oral pathologies (Abranches et al., 2011; Ikeda et al., 2023; Nakano et al., 2011; Nomura et al., 2013; Nomura et al., 2020).

Alongside these proteins involved in adhesion mechanisms, the role of the rhamnose glucose polysaccharide (RGP) clearly stands out as a key element in the virulence of *Streptococcus mutans*. This complex polymer, considered to be the equivalent of wall teichoic acids (WTA), may have a more important role than that of bacterial cell wall biogenesis. Roles such as phage receptor or immune response escape are attributed to it (Guerin et al., 2022; Mistou et al., 2016). This RGP is the basis for the classification of *S. mutans* into 4 different serotypes: c, e, f and k. Serotyping is a way of rapidly identifying any different strains of *S. mutans* that an individual may have in his or her oral cavity. This is of particular interest as some authors have shown that the concomitant presence of several serotypes increases the risk of caries (Seki et al., 2006; Shibata et al., 2003).

As previously stated, caries risk is increased when the diet is rich in refined sugars or when oral hygiene is deficient. These two factors are particularly common in Parkinson’s disease sufferers. Indeed, several studies agree that these subjects have a higher appetite for sugar, combined with impaired motor skills (Haas et al., 2022; Palavra et al., 2021; van Stiphout et al., 2018; Vanbellingen et al., 2011). As the presence and type of the above-mentioned adhesins are associated with an increase in comorbidities, understanding the distribution and prevalence of these adhesins is of particular interest in a fragile population.

Thus, in a population particularly susceptible to caries (the probability of finding *Streptococcus mutans* being higher), our work aimed to study the diversity of *S. mutans* strains and describe the distribution of the molecules of sucrose-independent adhesion.

## Materiels and methods

### Sample collection

All tested strains were isolated from saliva samples of patients included in the clinical trial: Preliminary Study of the Quality of the Oral Flora in Parkinson’s Patients (PARKIDENT, ClinicalTrials.gov Identifier: NCT03827551). Briefly, PARKIDENT was a prospective interventional clinical study whose main objective was to determine the impact of oral hygiene guidance on the oral health status of patients with Parkinson disease. Saliva samples were collected before and after the implementation of the hygiene measures, within a period of 6 months. The research protocol was approved by a French regional ethics committee (approval number: Eudract N° 2018-A02773-52), and all the participants signed a written informed consent form.

Dilution of the collected salivary samples were cultured on Mitis Salivarius Agar (Fischer Scientific, Illkirch, France) supplemented with potassium tellurite, bacitracin and kanamycin (Sigma-Aldrich, St Quentin Fallavier, France) for the streptococci. After examination of the colonies, gram staining was performed, followed by PCR from Oho et al. (Oho et al., 2000) and MALDI-TOF identification as previously described (Jussiaux et al., 2021).

All identified strains were kept frozen at -80°C.

### DNA extraction

DNA was isolated from each bacterial strain by using GenElute™ Bacterial Genomic DNA kit (Sigma Aldrich, France) according to the manufacturer’s supplied protocol.

### PCR Analysis

PCR analysis for the four serotypes and for collagen-binding proteins (Cnm, Cbm) were performed using multiplex PCR. Conditions used included an initial denaturation step of 3 min at 95°C, followed by 30 cycles of denaturation at 95° C for 30 s, primer annealing at 50°C for 45 s and extension at 72°C for 60 s. A final extension step was carried out for 2 min. All the PCR amplification products were examined using 1% agarose gel electrophoresis. A listing of all primers used in this study can be found in table 1 (Nakano et al., 2004; Nomura et al., 2012; Oho et al., 2000; Shibata et al., 2003).

**Table 1:**
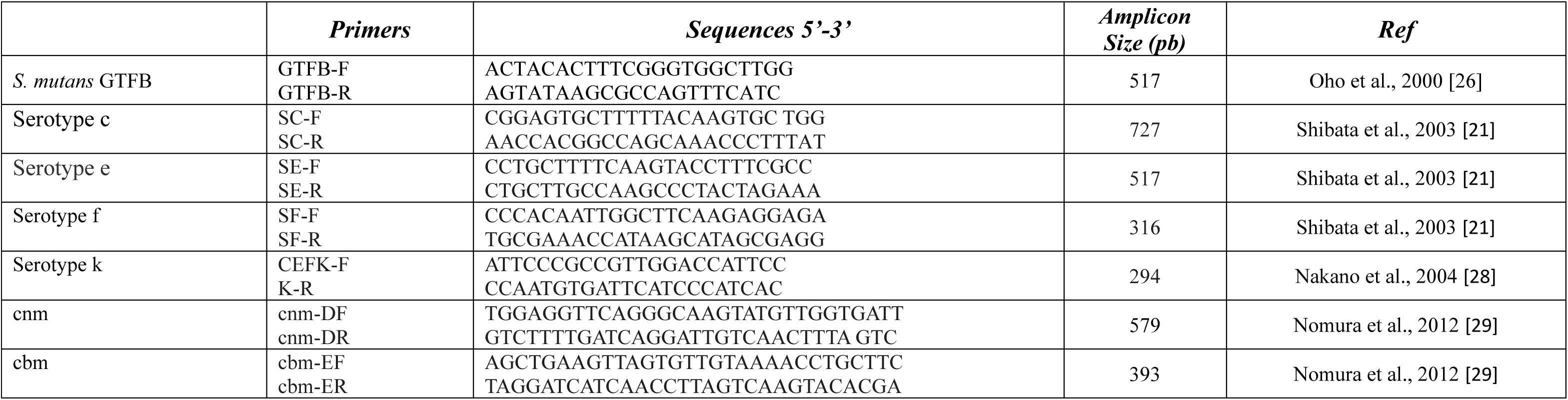
Primers used in this study

Control *S. mutans* strains used for PCR analysis were MT8148 (serotype c), NN2087 (serotype e), NN2117 (serotype f) and SA31 (serotype k, cbm+, cnm+).

### High-throughput sequencing

Whole genome sequencing of *S. mutans* strains was performed at the Genome-Transcriptome platform in Bordeaux (see table S1 for accession number) (Ziane-Casenave et al., 2023). DNA libraries were prepared using the Nextera XT DNA library preparation kit (Illumina, San Diego, CA). DNA were submitted to the Genome-Transcriptome Facility of Bordeaux for library preparation (QIAseq FX DNA; Qiagen, Courtaboeuf, France) and whole-genome sequencing (Illumina MiSeq V3); producing 2 × 300 bp paired-end reads. Sequences were trimmed using Trimmomatic v0.39 (Bolger et al., 2014) and assembled using Skesa v2.4.0 (Souvorov et al., 2018). The annotation with NCBI Prokaryotic Genome Annotation Pipeline (PGAP) v6.2 (Tatusova et al., 2016).

### Bioinformatics analysis

The computer analysis was carried out using the RAST® (Rapid Annotations using Subsystems Technology) and BLAST® (Basic Local Alignment Search Tool) software (Altschul et al., 1990; Aziz et al., 2008; Brettin et al., 2015; Overbeek et al., 2014). The raw data were loaded on RAST®, an automatic annotation server that which allows the assembly of raw sequences. The sequences of interest of the *S. mutans* genomes available on the databases (NCBI) were used on BLAST as queries to identify serotypes and collagen binding proteins.

Protein structure modeling was carried out using the web-based software Protter® (Omasits et al., 2014).

### Construction of a phylogenetic tree

All *S. mutans* sequences were collected and compared one by one with the ANI (Average Nucleotide Identity) algorithm. After obtaining the similarity matrix, an inverse conversion was performed to calculate the distances. These data are then used in MEGA (Molecular Evolutionary Genetics Analysis) to create the base of the tree. Finally, the data distance data obtained with MEGA are imported on the Interactive Tree Of Life website (https://itol.embl.de/) in order to create the final version of the tree (Kumar et al., 2018; Letunic & Bork, 2021).

## Results

### Strain information

Although 40 patients were included in the PARKIDENT clinical trial, it was possible to isolate *Streptococcus mutans* strains from the salivary swabs of only 24 patients. The 44 strains collected (shown in table 2) came from patients with active carious lesions as well as from patients without ongoing carious disease. 24 strains were isolated from samples taken at the inclusion visit and 20 at the follow-up visit.

**Table 2.** Strain information. Details of the source patient (anonymized), sampling visit, serotype, SpaP type and presence or absence of the *Cnm* gene are provided.

| Patient | Active caries status | Strain |  | Serotype | SpaP type | Cnm |
| --- | --- | --- | --- | --- | --- | --- |
|  |  | Inclusion | Follow-up |  |  |  |
| 01BG | Yes | 01BG6 |  | c | B | - |
|  | No |  | 01BGb2 | c | B | - |
| 02BD | No |  | 02BDb3 | c | B | +<br>(truncated) |
| 03GM | No | 03MG7 |  | c | A | - |
|  | No |  | 03GMb4 | c | A | - |
| 04BF | Yes | 04BF1 |  | e | B | - |
|  | Yes |  | 04BFb1 | e | B | - |
| 07TF | No | 07TF2 |  | c | B | - |
|  | No |  | 07TFb2 | c | B | - |
| 08CA | No | 08CA2 |  | c | B | - |
|  | No | 08CA3 |  | c | B | - |
|  | Yes |  | 08CAb2 | c | B | - |
| 09DC | Yes | 09DC3 |  | c | B | +<br>(truncated) |
|  | No |  | 09DCb1 | c | A | - |
| 11MM | Yes | 11MM2 |  | k | A | - |
|  | No |  | 11MMb3 | c | B | +<br>(truncated) |
| 12RB3 | Yes | 12RB3 |  | c | B | - |
| 14BP | Yes | 14BP2 |  | k | A | - |
|  | Yes | 14BP3 |  | c | A | - |
| 15LT | Yes | 15LT2 |  | c | A | - |
| 16SP | Yes | 16SP2 |  | c | B | + (shorter) |
| 17PM | No | 17PM2 |  | c | B | - |
|  | No |  | 17PMb4 | c | B | + (shorter) |
| 18MJ | No | 18MJL1 |  | c | A | - |
|  | Yes |  | 18MJLb4 | c | B | - |
| 19CL | Yes |  | 19CLb3 | c | B | - |
| 20QP | Yes | 20QP1 |  | c | A | - |
|  | Yes | 20QP2 |  | c | A | - |
|  | Yes | 20QP3 |  | c | A | - |
| 22HM | Yes |  | 22HMb3 | e | A | - |
| 24CC | No | 24CC1 |  | c | Truncated | - |
|  | No |  | 24CCb1 | c | A | - |
|  | No |  | 24CCb2 | c | A | - |
| 25VC | Yes | 25VC3 |  | c | B | - |
|  | Yes |  | 25VCb1 | c | B | - |
|  | Yes |  | 25VCb3 | c | B | - |
| 26BM | Yes | 26BM3 |  | k | B | - |
| 33HA | No | 33HA1 |  | c | A | - |
| 34BR | Yes |  | 34BRb2 | e | B | - |
| 35DF | No | 35DF1 |  | c et k | A | - |
|  | No | 35DF2 |  | c | A | - |
|  | No |  | 35DFb1 | c | A | - |
| 37PJ | No |  | 37bPJ2 | c | A | - |
|  | No |  | 37bPJ3 | c | A | - |

### Serotype distribution of the strains

Multiplex PCR for serotyping was carried out on the 44 strains isolated in this study. The data obtained showed that over 80% of the strains were serotype c (shown in table 2). Due to the difficulty of differentiating serotype f from serotype k on agarose gel alone (data not shown), a further check was carried out using sequencing data. This analysis showed that none of the strains in this study was serotype f. In order to compare the serotyping data of the clinical isolates in this study with those already available, an analysis was also made of all the *Streptococcus mutans* genomes available in the databases of the National Center for Biotechnology Information (NCBI). The search carried out on July 12, 2022 found 352 strains, 332 of which could be grouped into one of the serotypes already described on the basis of specific peptide sequences (see figure 1). Of the 20 strains left to be characterized, 2 could not be classified because the level of identity with the rgpF-encoded protein was insufficient (less than 92.1%) to classify them as serotype k, while the other 18 had a double serotype. The predominant double serotype was c and k (over 83% of the 18 strains). Of the *S. mutans* clinical isolates in this study, only strain 35DF1 was found to a double serotype (c and k). Similarly to the clinical isolates in the PARKIDENT study, of the 352 *S. mutans* sequences found, the majority are serotype C (just over 80%).

**Figure 1:**
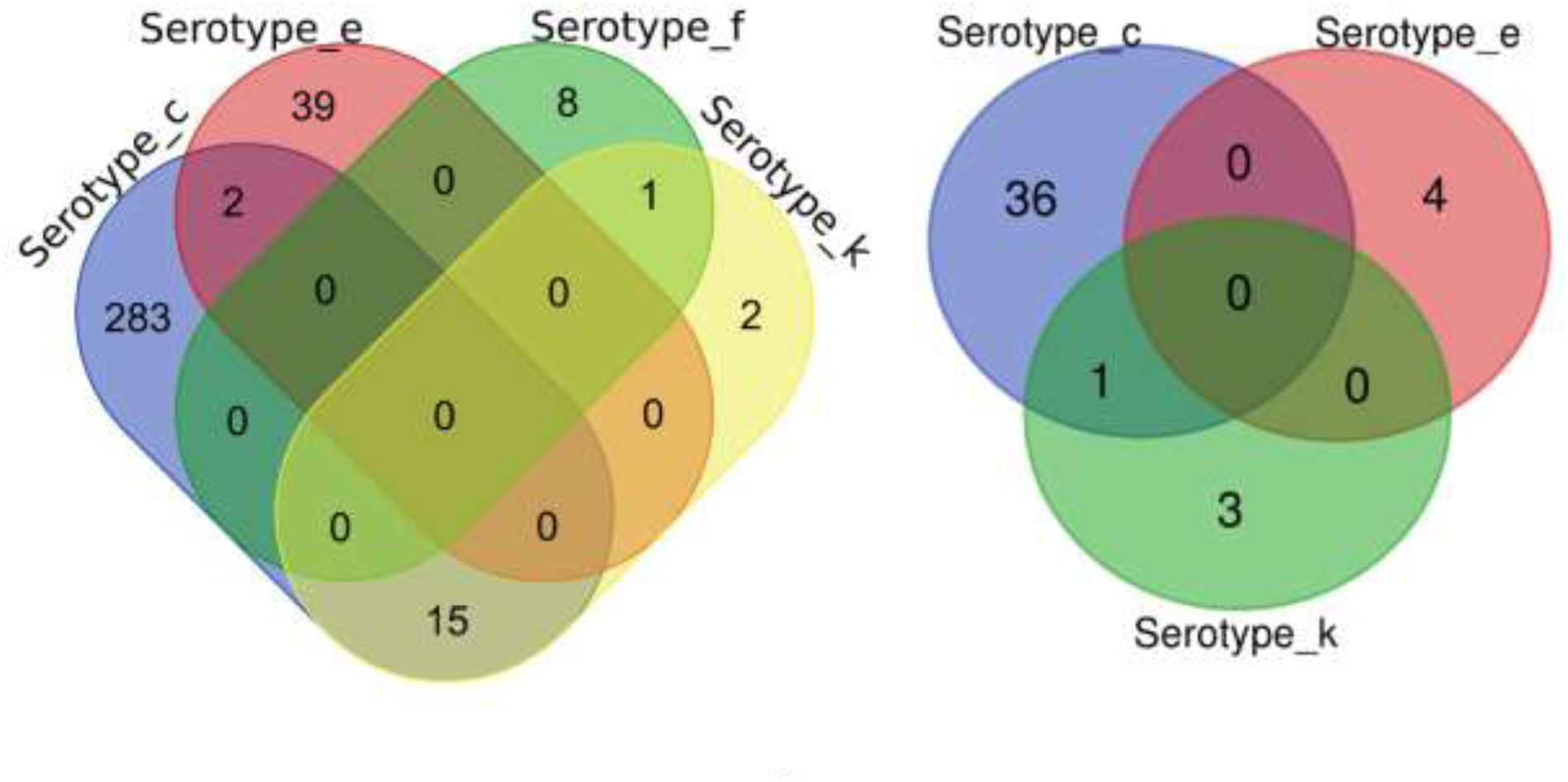
Serotype distribution of the 44 strains in this study: comparison with 350 strains in a public database. The Venn diagrams used to classify strains according to serotype were produced using protein sequences whose accession numbers are available in the Protein database on the National Center for Biotechnology Information (NCBI) website. The sequences WP_011074554.1 (rgpH gene product) for serotype c, WP_106006773.1 (ORF 3e gene product) for serotype e, WP_002270463.1 (ORF 2f gene product) for serotype f and BAL69517.1 (rgpF gene product) for serotype k. The Venn diagrams were constructed using the Bioinformatics & Evolutionary Genomics website (http://bioinformatics.psb.ugent.be/webtools/Venn/).

### SpaP type of the strains

No strains with a SpaP type C protein were detected in this study. The strains are either type A or type B, the distribution between these 2 types being rather homogeneous (Fig. 2, see accession number in table S2). SpaP protein sequences are relatively close, with less than 5% variability from the sequence available under the UniProtKB C9E3B4 reference (figure 2, see details in table S3). The protein sequence of strain 24CC1 is truncated, making it impossible to classify it in the phylogenetic tree (see figure 2).

**Figure 2:**
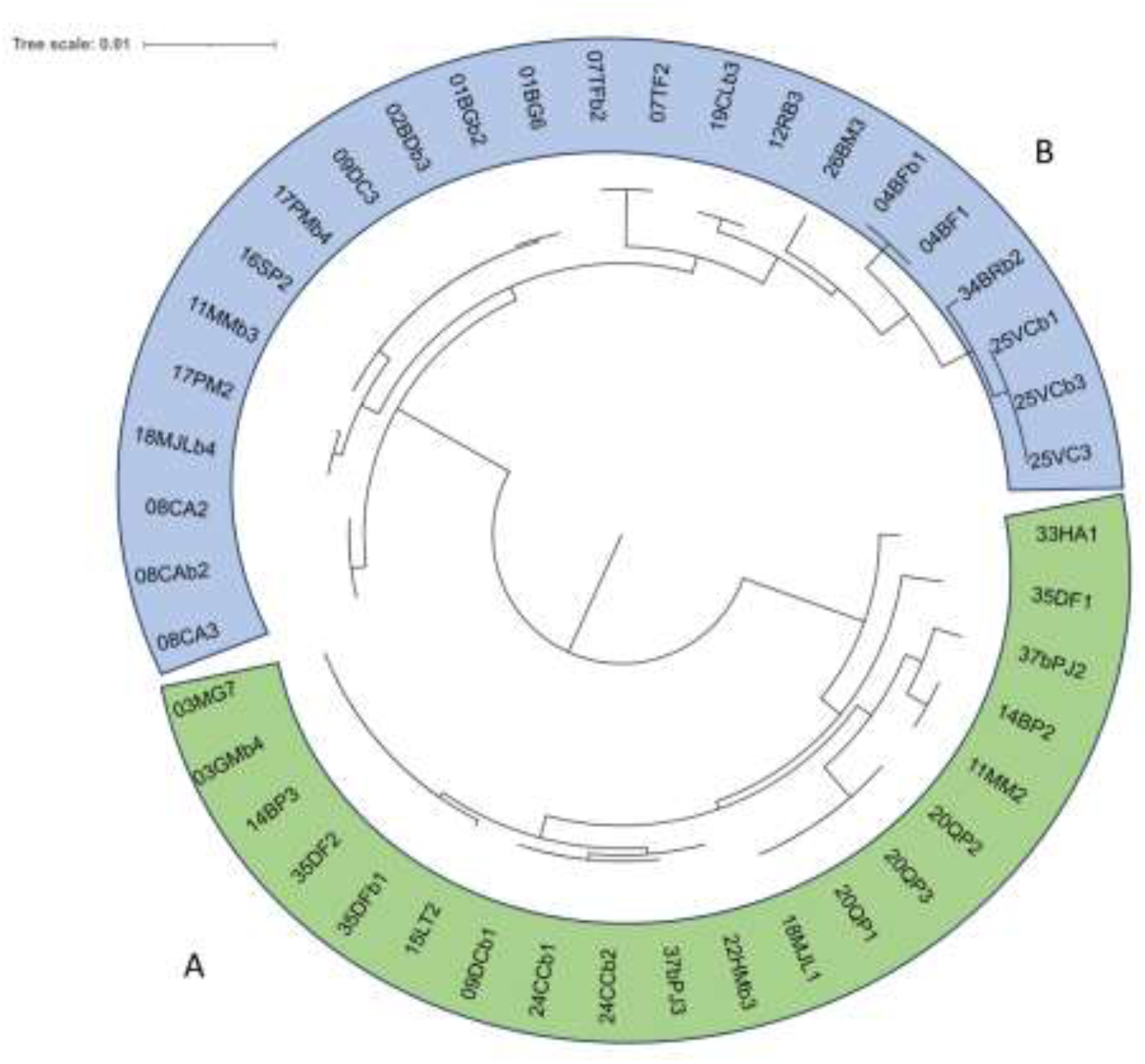
Phylogenetic tree showing SpaP diversity for each of the 44 strains in this study. The phylogenetic tree was constructed on the basis of the SpaP protein sequences for all strains (except strain 24CC1) and taking as reference the protein sequence available under accession number UniProtKB:C9E3B4. SpaP type A strains are grouped on the same side of the tree, the same is true for type B. Details of SpaP identity percentages for each strain with the reference are given in supplemental materials, in Table S3. In Table S2, the SpaP sequence accession numbers for each strain are given.

### Variability of WPA

The WPA protein gene is present in all strains. It also appears to be functional in all strains, since it is never truncated. Protein sequence identity is high, with less than 1% variability (figure 3, see details in table S3). The protein sequence used as a reference is available under accession number UniProtKB P11000.

**Figure 3:**
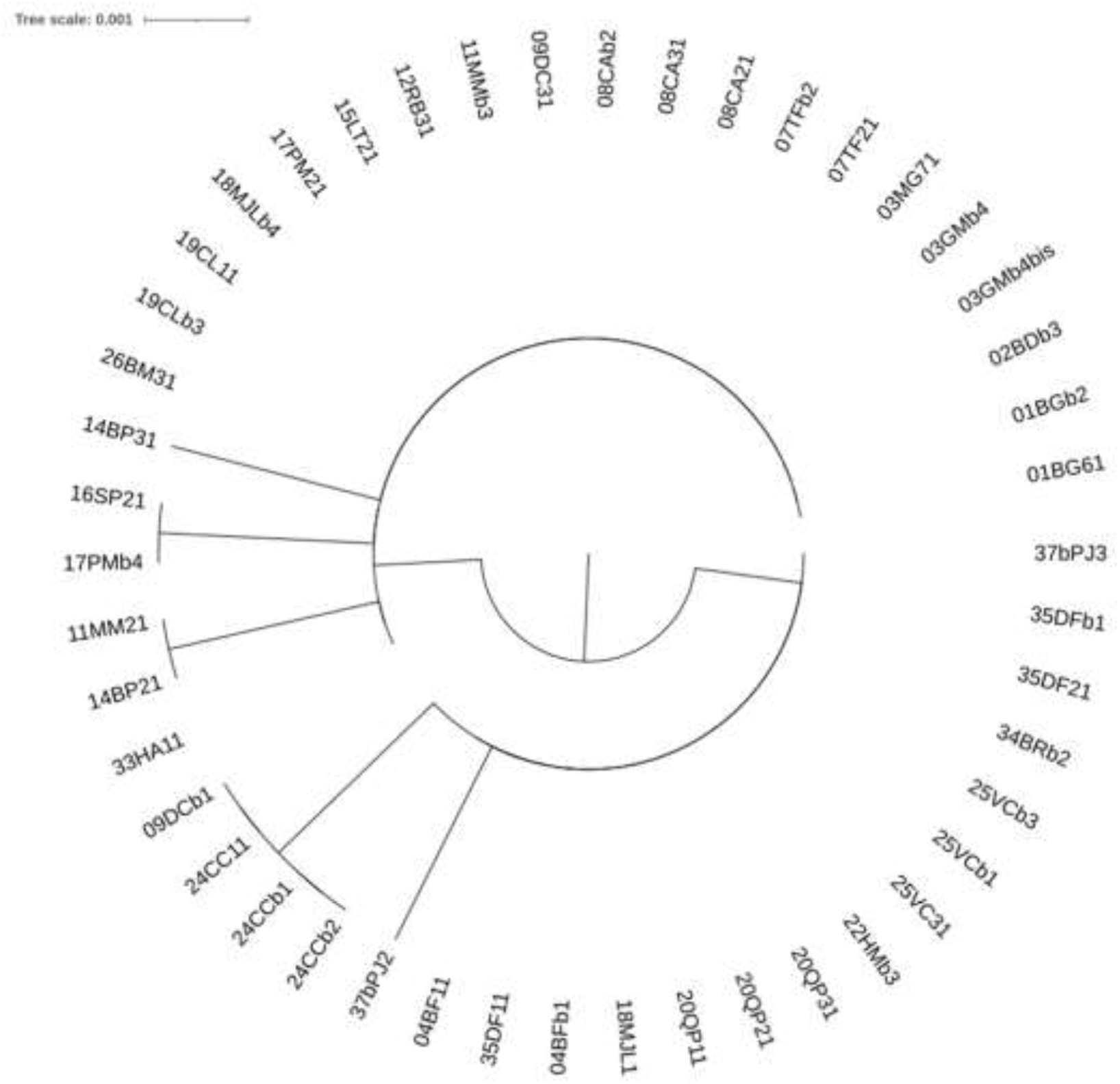
Phylogenetic tree showing WapA diversity for each of the 44 strains in this study. The phylogenetic tree was constructed on the basis of the WapA protein sequences for all strains, taking as reference the protein sequence available under UniProtKB accession number P11000. Details of the percent identity of WapA for each strain with the reference are given in in table S3.

### Cnm and Cbm adhesins

A rapid multiplex PCR search for the presence of the *cnm* and *cbm* genes showed that no strains possessed the *cbm* gene and only 5 strains the *cnm* gene. All Cnm-positive strains were found to be serotype c (shown in table 2). Sequencing data from these strains showed that three of them (02BDb3, 09DC3 and 11MMb3) carried a gene coding for a truncated protein. The Cnm protein sequences of strains 16SP2 and 17PMb4 were compared with the Cnm sequence of strain TW871 (available as UniProtKB C4B6T3). The proteins encoded by these two strains are 81 amino acids smaller than the reference Cnm protein (549 amino acids). The deleted portion corresponds to a repeat motif (figure 4).

**Figure 4:**
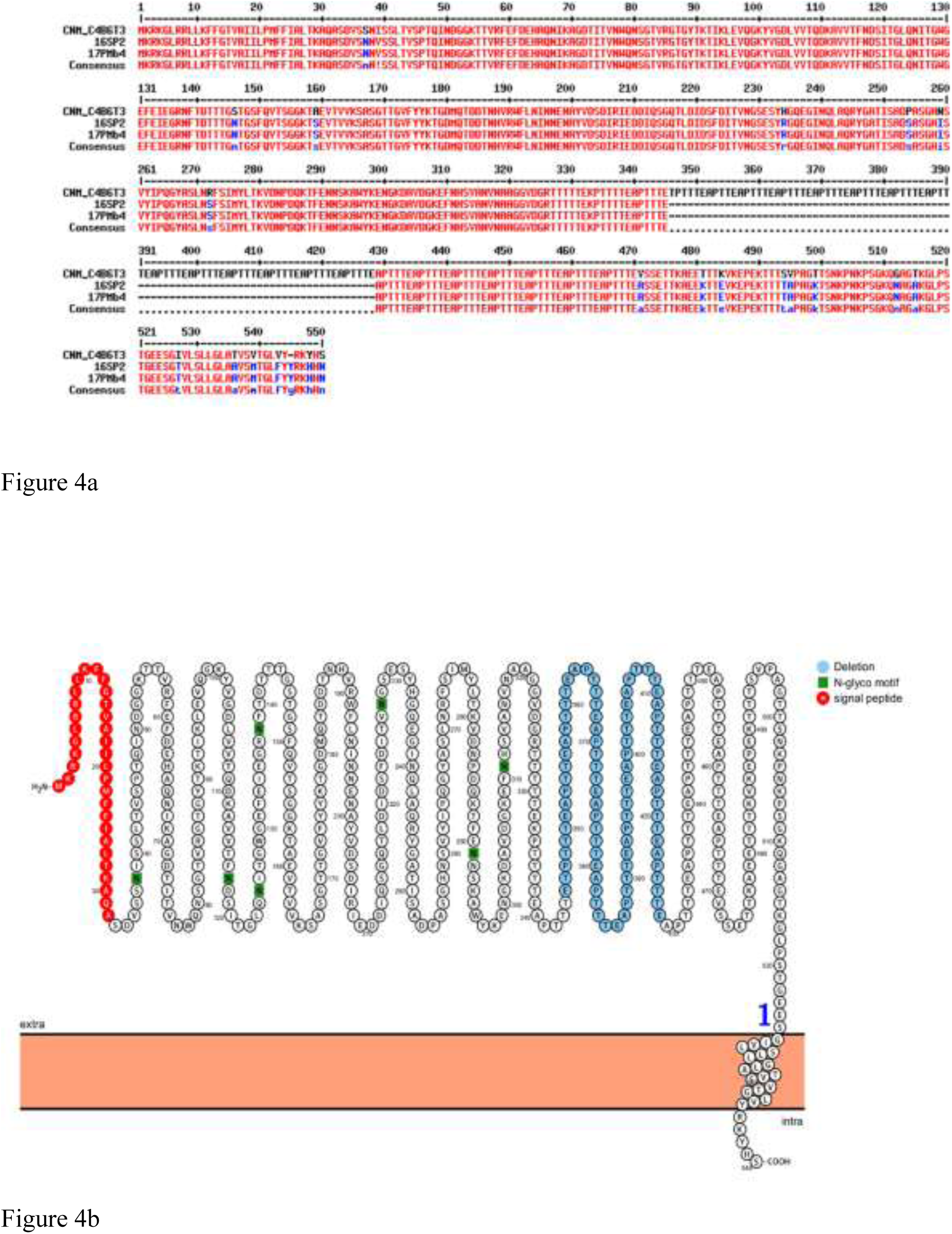
Exploring the diversity of functional Cnm proteins in this study. Peptide alignment of the Cnm sequences of the 2 strains (16SP2 and 17PMb4) with a non-truncated protein. The alignment was carried out with the reference sequence available under the UniProtKB accession number C4B6T3. Visualization of the predicted Cnm proteoform. The wall-addressing signal peptide is colored red, the deleted portion in the proteins from strains 16SP2 and 17PMb4 is blue, and the number 1 indicates the number of transmembrane domains.

## Discussion

*Streptococcus mutans* is an essential bacterium in the composition of dental biofilm. Since it is also involved in extra-oral pathologies, we consider its study and epidemiological follow-up in a given subject to be important, as do many authors. Indeed, strains presenting certain adhesion proteins (Cnm, Cbm) or a specific stereotype (serotype k) have been implicated in non-oral pathologies. Moreover, *S. mutans* remains strongly associated with caries experience, even if its presence alone at a given threshold is not sufficient to explain the presence of caries in all individuals.

In this study, of the 40 participants enrolled in the PARKIDENT study, salivary samples from only 24 subjects enabled *Streptococcus mutans* strains to be isolated. This recovery rate seems low compared with some published data (Momeni et al., 2019) but concordant with others (Esberg et al., 2017). The relevance of the saliva sample could be questioned, but recent studies have reaffirmed that the salivary recovery of *S. mutans* is comparable to that of dental plaque (Nguyen et al., 2022). On the other hand, although the selective medium used in this study remains a reference, recent data suggest that *S. mutans* isolation could be optimized with other media (Saravia et al., 2020; Singla et al., 2016).

The pathogenicity of *Streptococcus mutans* is largely explained by the different virulence factors present in this species. Deciphering the complexity and exploring the diversity of virulence factors in bacterial pathogens are essential prerequisites for treating the infectious diseases for which these pathogens are responsible. As adhesion is an essential factor in bacterial virulence, here we focused on a range of adhesins already well identified in *S. mutans*, but also on the species’ rhamnose glucose polysaccharide (RGP), which has been shown to play a critical role in the bacterium’s pathogenesis (Kovacs et al., 2019). The presence of this RGP on the bacterial surface enables it to be classified into 4 different serotypes: c, e, f and k. In this work, based on sequencing results, we report for the first time, to our knowledge, epidemiological data on *S. mutans* in an adult French population, as well as serotyping data on over 300 strains whose genomes are publicly available. One of the most surprising results of this study is the fact that some strains show characteristics that allow them to be classified in 2 different serotypes. Momeni et al. had already made this observation using multiplex PCR, justifying it by a lack of primer specificity for serotype c detection and suggesting that new primers for serotype c be designed (Momeni et al., 2019). Here, unlike the only PCR-based approach which targets the nucleotide sequence of target genes (rgpH, ORF3e, ORF2f, rgpF) for serotype determination (serotype c, e, f and k respectively), we also used the protein sequence of the product of these genes. This did not allow us to completely rule out the hypothesis of strains possessing a double serotype, i.e. two different types of RGP on their surface. The existence of a double serotype has yet to be proven with further functional research. However, on the one hand, authors report that data concerning the ability of RGP-expressing strains to vary its expression, composition and length are unclear (Mistou et al., 2016), and on the other, some strains remain untyped, especially in subjects with extra-oral pathologies in which *S. mutans* is involved (Nakano & Ooshima, 2009). This could be in favor of a diversity of RGP polymers on the same strain. Precise serotype determination seems important, as some authors have shown that serotype k strains are more likely to be linked to persistent bacteremia: the absence of glucose on the rhamnose backbone would not favor phagocytosis (Nagata et al., 2006). Studies have also shown that glucose side chain of RGP is also important in *S. mutans* for bacteriophage adhesion (Shibata et al., 2009). Similarly to many other studies, in our study the most frequently found serotype is serotype c (Hirasawa & Takada, 2003; Nakano & Ooshima, 2009). The most plausible hypothesis is that serotype c is the ancestral form from which the other serotypes were derived (Maruyama et al., 2009).

Three types of SpaP have been described in the literature (Esberg et al., 2017). In our study, only two types were found: type A and type B. This seems to be in line with other authors (Yang et al., 2019). The variability of SpaP, although low, is to be compared with that of its ligand: salivary agglutinin (SAG)/DMBT-1 (Alharbi et al., 2022). Salivary agglutinin is a pattern recognition receptor for many bacteria, including *S. mutans* (Ligtenberg et al., 2010). Polley et al. have shown that there is a co-evolution between SAG and agriculture, and in particular the increased consumption of refined sugars (Polley et al., 2015). We can therefore hypothesize that *S. mutans* strains, via SpaP modifications, have also evolved with our diet. The findings of a study by Esberg et al., which consider that strains with a type B SpaP could be associated with a higher caries risk seem to support this hypothesis (Esberg et al., 2017). A contrario, we could imagine that a strain with a truncated SpaP (like strain 24CC1 here) could be deficient in its initial adhesion but also in biofilm formation since it has been shown that SpaP also binds, for example, with the RadD of *Fusobacterium nucleatum ssp polymorphum* (Guo et al., 2017).

In contrast to SpaP, there are fewer data in the literature concerning WapA (also known as antigen III). Its importance for S. mutans seems to be greater than that attributed to it in the scientific literature. Indeed, in our study, this adhesin shows very low sequence variability (less than 1%) and appears to be functional in all strains (no premature stop codon). The conservation of this protein in all strains tends to confirm its importance. However, some authors underline its importance in cell chain and biofilm formation (Li et al., 2015; Zhu et al., 2006).

The other two adhesins investigated in this study are collagen-binding proteins (CBP). Cnm and Cbm are found inconsistently in *S. mutans*, with studies reporting frequencies of around 20% and 5% respectively. The results here are fairly concordant concerning the low distribution of Cbm in *S. mutans*, since none of our strains had the gene encoding this protein. The *cnm* gene is more frequently found, with 5 strains displaying the gene, but 3 of them having a gene coding for a truncated protein. The two remaining genes code for a smaller Cnm protein than the one used as a reference (C4B6T3), but this seems to have little impact on the functionality of the encoded protein. The signal peptide is retained, as is the collagen-binding domain (CBD) and the LPXTG motif that enables adhesin to be addressed at the bacterial cell surface. The deleted amino acids appear to be involved only in beta-sheet formation. The collagen-binding function of Cnm (monomeric form) may be fully functional, while its polymerization, which enables amyloid aggregation, may be disrupted by the deletion affecting the area involved in beta-sheet formation. Amyloid aggregation is also a capacity that has been demonstrated in other *S. mutans* proteins: the adhesins SpaP (notably its derivative portion C123) and WapA (via its truncated product AgA) but also Smu_63c (a negative regulator of biofilm cell density and genetic competence) (Besingi et al., 2017). The importance of these functional amyloids in the constitution of biofilms means that these proteins are now targeted by anti-biofilm therapeutics (Matilla-Cuenca et al., 2021; Paranjapye & Daggett, 2018). It should be noted that all strains carrying the Cnm gene in this study are serotype c confirming recent observations made on the subject (Momeni et al., 2019; Nakano et al., 2010).

This variability in the presence of virulence factors in *S. mutans* confirms the findings of other authors who consider that the S. mutans genome is an open genome with a relatively small core genome (Bedoya-Correa et al., 2019; Meng et al., 2017). The accessory genome therefore plays a more important role, which may explain the difference in pathogenicity between strains, but above all underlines its adaptability and co-evolution with its host.

## Conclusion

The study of 44 strains of *Streptococcus mutans* from the oral cavity of this population of people suffering from Parkinson’s disease did not differ significantly from the characteristics found in most other populations studied to date. Serotype c is the most prevalent, and there is little peptide variability in the main proteins involved in non-sucrose-mediated adhesion (SpaP and WapA). The collagen-binding proteins Cnm and Cbm are rarely (for Cnm) or not at all (for Cbm) found in these clinical isolates. Further studies are nevertheless needed to investigate in greater detail the distribution of *S. mutans* strains in these specific populations affected by Parkinson’s disease, but also in the French population in general, especially as *Streptococcus mutans* is increasingly implicated in extra-oral pathologies.

## Acknowledgments

We warmly acknowledge Martine Saint-Marc for the technical assistance in the laboratory. We would also like to express our deepest gratitude to Prof. Kazuhiko Nakano and Prof. Ryota Nomura for their contribution to this study by kindly sending us the control strains.

Our last words are dedicated to the memory of Dr Marie-Cécile Badet, who recently passed away and contributed to the collection of the strains.

## Statement of Ethics

The research protocol was reviewed and approved by a French regional ethics committee, the Comité de Protection des Personnes (CPP) Sud-Est III, approval number: Eudract N° 2018- A02773-52.

All participants signed a written informed consent.

## Author Contributions

LD: formal analysis, writing - original draft; OC: data curation, formal analysis conceptualization, and writing - review and editing; JS: project administration, conceptualization, methodology, funding acquisition, investigation, formal analysis, validation, writing - original draft, and writing - review and editing

## Data Availability Statement

All data generated or analyzed during this study are included in this article and its supplementary material files. Further enquiries can be directed to the corresponding author.

---

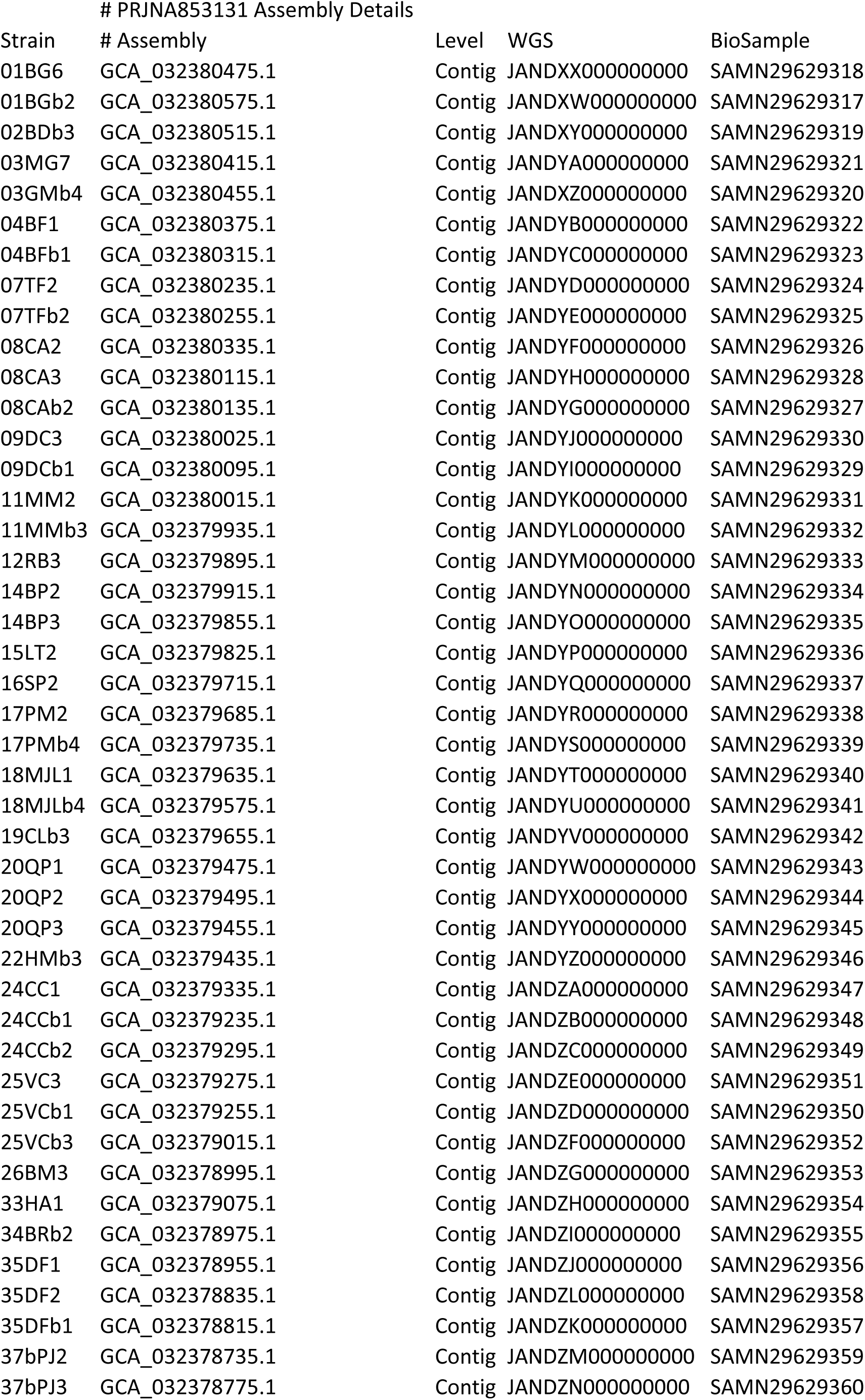
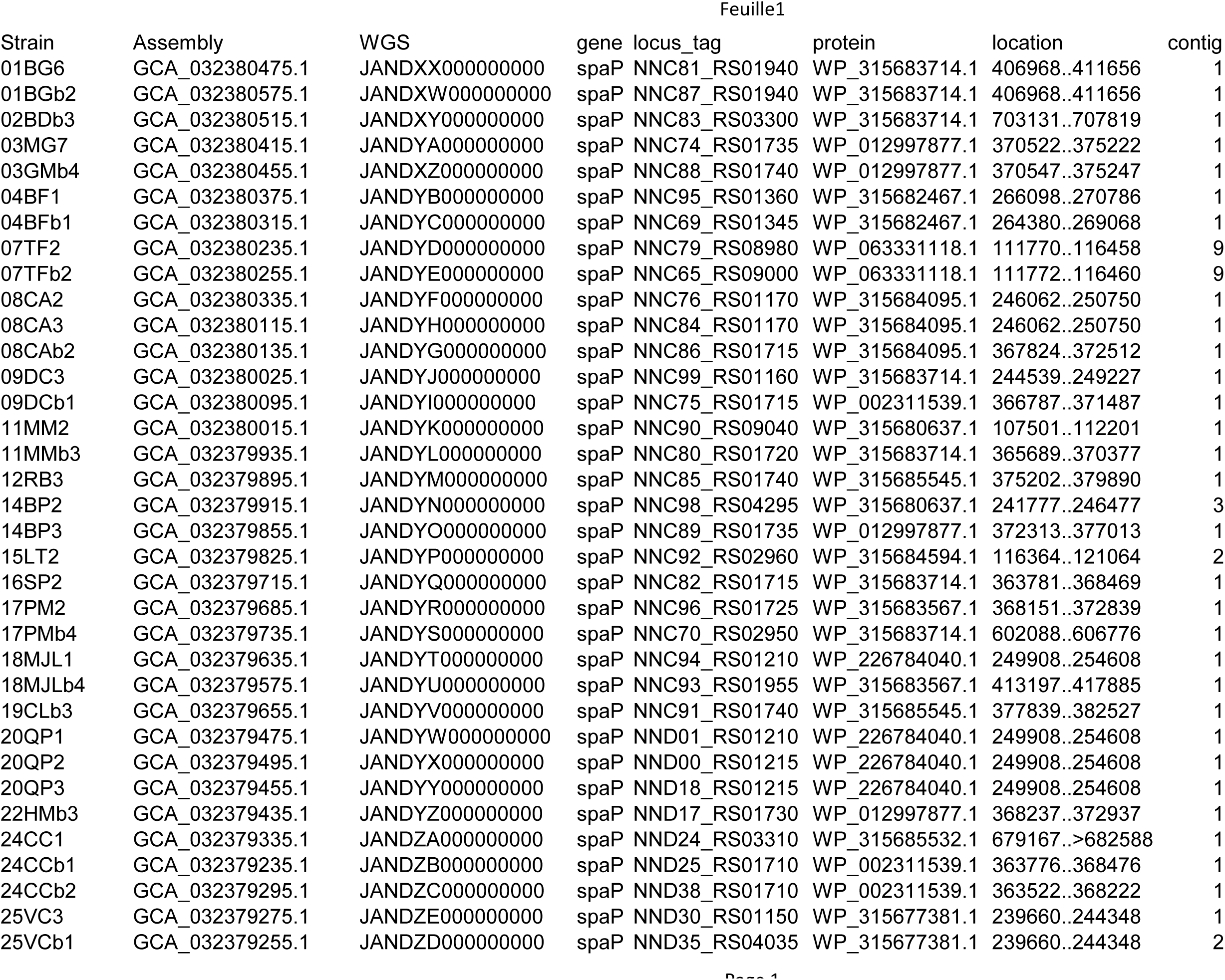
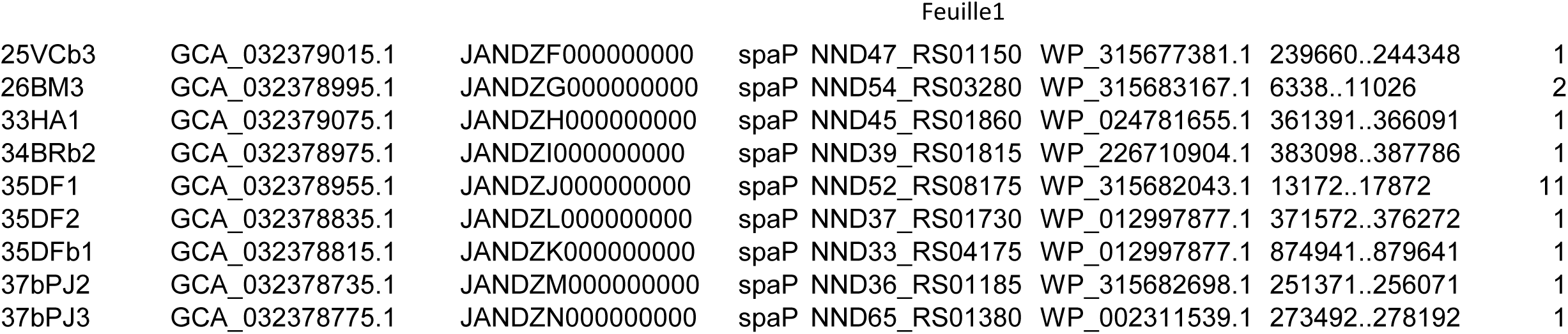
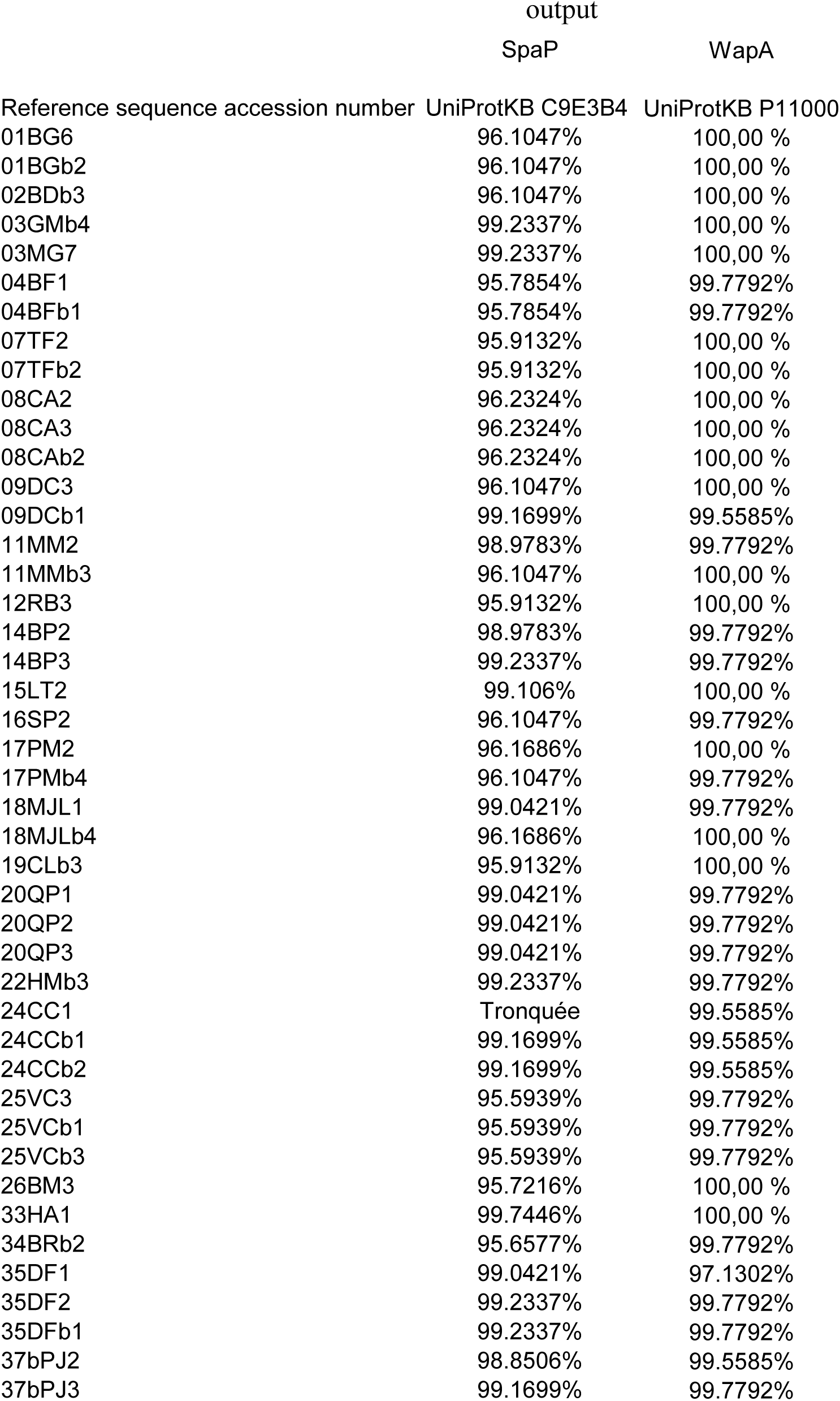

